# Coherent Structures in Active Flows on Dynamic Surfaces

**DOI:** 10.1101/2025.05.23.655805

**Authors:** Sreejith Santhosh, Cuncheng Zhu, Blase Fencil, Mattia Serra

**Affiliations:** Department of Physics, University of California, San Diego, California 92093, USA; Department of Mechanical Engineering, University of California, San Diego, California 92093, USA

## Abstract

Coherent structures—flow features that organize material transport and deformation—are central to analyzing complex flows in fluids, plasmas, and active matter. Yet, identifying such structures on dynamic surfaces remains an open challenge, limiting their application to many living and synthetic systems. Here, we introduce a geometric framework to extract Lagrangian and Eulerian coherent structures from velocity data on arbitrarily shaped, time-evolving surfaces. Our method operates directly on triangulated meshes, avoiding global parametrizations while preserving objectivity and robustness to noise. Applying this framework to active nematic vesicles, collectively migrating epithelial spheroids, and beating zebrafish hearts, we uncover hidden transport barriers and Lagrangian deformation patterns—such as dynamic attractors, repellers, isotropic and anisotropic strain—missed by conventional Eulerian analyses. This approach offers a new perspective on soft and living matter, revealing how geometry and activity can be harnessed to program synthetic materials, and how Lagrangian strain and principal deformation directions can help uncover mechanosensitive processes and directional cues in morphogenesis.

## 1 Introduction

Active matter flows on curved surfaces are ubiquitous in biological systems, spanning morphogenesis [1–3], cytoskeletal remodeling [4, 5], in vitro epithelial spheroids [6–9], and synthetic active nematics [10, 11]. Recent advances in live imaging and computation have enabled the reconstruction of such flows across systems ranging from developing embryos to engineered active nematics [12–16]. These flows are spatially heterogeneous and temporally unsteady, driven by internal activity and influenced by surface curvature and topology [9, 11, 17]. While kinematic velocity or trajectory data becomes increasingly available, its analysis remains challenging, yet essential for controlling material transport or discovering the underlying biophysical mechanisms.

Lagrangian and Eulerian Coherent Structures—denoted together as Coherent Structures (CSs)—provide a reduced representation of complex trajectories, organizing material transport and deformation [18–21]. CSs are typically objective [22]—i.e., invariant to the choice of coordinate system adopted to describe motion—and robust to noise, hence ideal for analyzing and comparing experiments. CSs have been broadly applied in science and engineering [23], including atmospheric and oceanic flows [24–27], turbulence [28, 29], plasmas [30], cardiovascular flows [31], bioinspired flows [32], as well as to study transport of planktons in jellyfish feeding [33] and dispersion of plant pathogens [34]. In active nematics, recent works elucidated the connection between CSs and topological defect motion, revealing the interplay between flow structure and nematic order [35–37]. Similarly, Lagrangian CSs revealed new cumulative deformation patterns and dynamic boundaries in multicellular flows–robust to noise and experimental variability–while remaining undetected by Eulerian quantities [38]. These structures provided new insights into morphogenetic mechanisms [39, 40], pattern formation [41], and cell fate bifurcations [42, 43]. While existing CSs techniques cover general two- and three-dimensional flows, flows defined by sparse, noisy trajectories [44–47] or flows on static surfaces [38, 48], they remain unavailable for flows on dynamic surfaces—ubiquitous in natural and synthetic active systems.

Here, we introduce a theoretical and computational framework to identify Lagrangian and Eulerian CSs in flows on arbitrary dynamic surfaces. Our method estimates deformation tensors from velocity data defined on triangulated meshes, eliminating the need for global parameterizations and extending the CS framework to evolving surfaces. We apply this framework to nematic flows on deforming vesicles, collective migration in pancreatic spheroids, and cardiac tissue deformations in embryonic zebrafish hearts—revealing undocumented dynamic attractors, repellers and deformation patterns. These CSs and directional patterns of maximal stretch or contraction—absent in instantaneous velocity or nematic fields (Fig. 1)—can help uncover emergent mechanical properties and mechanotransduction mechanisms in living and synthetic active matter using only available kinematic data. We provide a brief tutorial and MATLAB and Python software (link) to facilitate use by the biological and soft matter communities.

**Figure 1:**
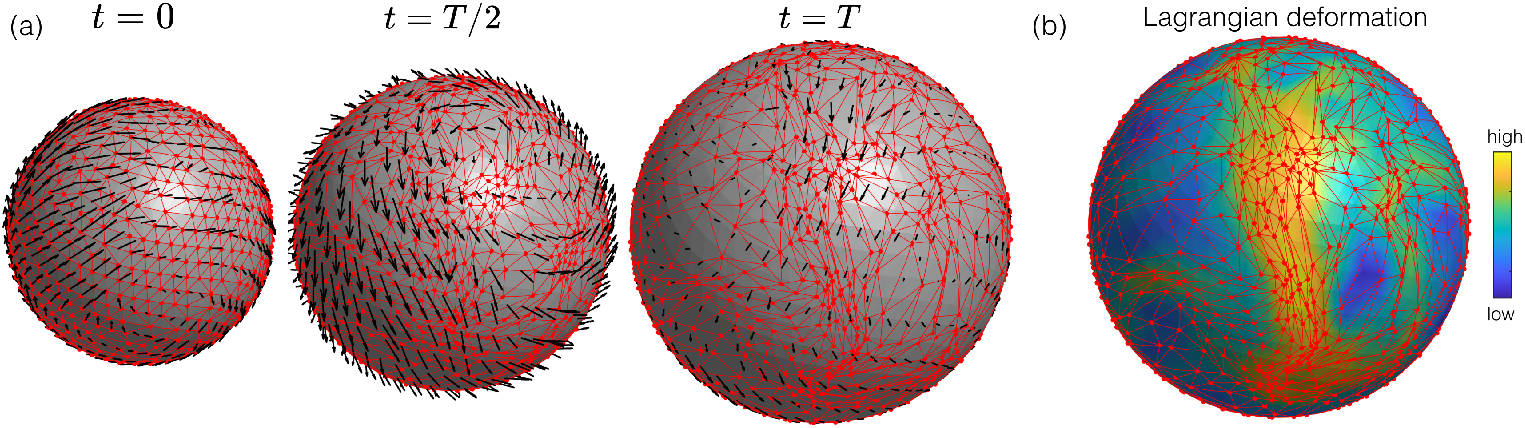
Lagrangian Coherent Structures provide the geometric skeleton of complex flow fields. (a) Snapshots of unsteady velocity fields (black) tangent to a growing spherical surface and the corresponding Lagrangian deformation grid (red). (b) Example of Lagrangian Coherent Structures marking regions of maximal material contraction (or attraction) over [0, *T*]. High deformation regions (yellow) are confirmed by the deforming grid while undetected by Eulerian velocities snapshots.

## 2 Results

### 2.1 Lagrangian Coherent Structures on dynamic surfaces

Flows on dynamic surfaces are represented by a three-dimensional velocity field **v**(**x**, *t*) defined on a time-dependent two-dimensional surface ℳ(*t*). ℳ(*t*) is typically provided in the form of a triangulated mesh (a set of connected points in 3D) on which **v**(**x**, *t*) is measured or simulated. Throughout, we denote vectors and matrices in bold and scalars in normal font. To study material transport and deformations of these flows over a time-interval [*t*_0_, *t*], we define the flow (or trajectory) map

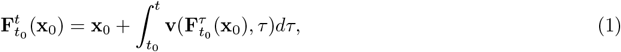

which maps the initial position (*x*_0_, *y*_0_, *z*_0_) = **x**_0_ ∈ ℳ (*t*_0_) of a material particle to its final position 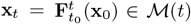 (Fig. 2a-b). The deformation experienced by a material patch in the neighborhood of **x**_0_ is encoded in

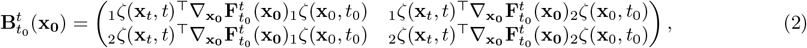

where [_1_*ζ*(**x**, *t*), _2_*ζ*(**x**, *t*)] are orthonormal basis vectors for the tangent space *T*_**x**_ℳ(*t*) (Fig. 2c), 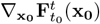 is the flow map gradient and ^⊤^ denotes vector transposition (SI Sec. 1 for derivation). To visualize 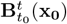, consider a material fiber 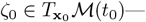 a vector connecting two infinitesimally close material particles in the neighborhood of **x**_0_, expressed in the basis vectors [_1_*ζ*(**x**_0_, *t*_0_), _2_*ζ*(**x**_0_, *t*_0_)]. *ζ*_0_ moves and deforms onto the material fiber 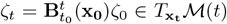, represented in the corresponding, orthonormal basis [_1_*ζ*(**x**_*t*_, *t*), _2_*ζ*(**x**_*t*_, *t*)] (Fig. 2c, SI Sec. 1). The maximum stretching experienced by any material fiber 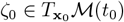 at **x**_0_ corresponds to the largest singular value 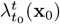 of 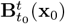, from which one can compute the largest Finite-Time Lyapunov Exponent (FTLE)

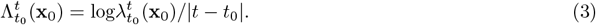

**Figure 2:**
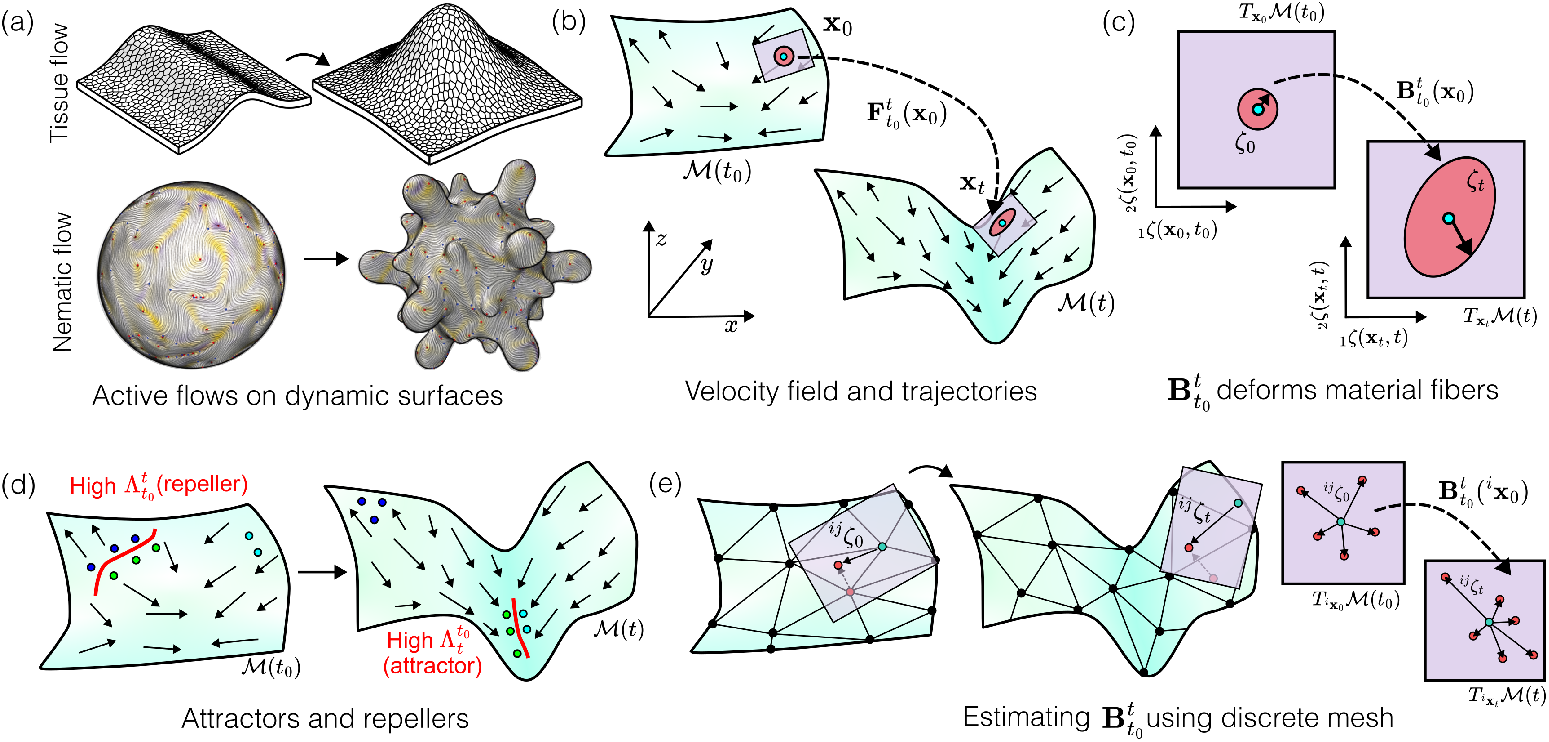
Lagrangian deformation and Coherent Structures of flows on dynamic surfaces. (a) Schematic tissue and nematic active flows on dynamic surfaces. (b) Flows are represented by three-dimensional velocities **v**(**x**, *t*) on evolving two-dimensional surfaces ℳ(*t*). The flow (or trajectory) map 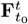 maps the initial position **x**_0_ of a material particle (cyan dots) to its final position **x**_*t*_. (c) The material deformation experienced in the neighborhood of **x**_0_ (purple patch) is encoded in 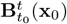 (eq. 2), and [_1_*ζ*(**x**, *t*), _2_*ζ*(**x**, *t*)] ∈ *T*_**x**_ ℳ (*t*) represents a local orthonormal basis for the tangent space at **x** ∈ ℳ (*t*) (pink square). (d) Material particles (blue, green) on either side of a repeller—marked by high values of 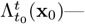 maximally separate during [*t*_0_, *t*]. By contrast, attractors—marked by high values of 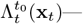 locate final positions where initially far away particles (green, cyan) will maximally converge during 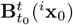 encodes deformations of material fibers in the neighborhood (or tangent plane) of ^*i*^**x**_0_, and can be estimated for each mesh node *i*, using the relation 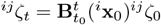 for all neighboring nodes *j* (SI Sec. 2).

The material fiber (i.e. the local direction) *ζ*_0_ at **x**_0_ which undergoes maximal stretching corresponds to the largest singular vector 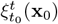 of 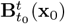. Similarly, one can quantify the isotropic Lagrangian deformation by

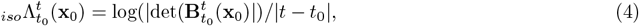

where 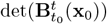 represents the ratio of the final (time *t*) to the initial (*t*_0_) area of an infinitesimal material patch starting at **x**_0_ (Fig. 2b-c). Regions of high 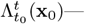 i.e., forward-time FTLE or _*f*_ Λ(**x**_0_)—mark initial domains where initially close particles will maximally separate in forward time during [*t*_0_, *t*], locating *repelling LCS* [18, 19, 49], hereafter called *repellers*. Similarly, the backward FTLE 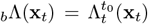 field— computable by integrating trajectories in backward time from *t* to *t*_0_—is a scalar field over the final positions **x**_*t*_, whose high value marks the location of *attracting LCSs*, hereafter *attractors* (Fig. 2d). Therefore, given a time interval [*t*_0_, *t*], repellers are visualized at the initial positions (marked by high values of _*f*_ Λ(**x**_0_)) while attractors at the final position (marked by high values of _*b*_Λ(**x**_*t*_)). Together, _*f*_ Λ(**x**_0_), _*b*_Λ(**x**_*t*_) and _*iso*_Λ fully characterize Lagrangian isotropic and anisotropic deformation. While there are multiple definitions and types of LCSs [19,21,50–54]—some of which directly computable from 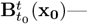 here we focus on the most common one– attractors and repellers based on FTLE–owing to its computational simplicity and straightforward visualization.

In simple cases, ℳ(*t*) can be described via a parametrization **x** = **r**(**p**), where **p** denotes the parameters. In this case, the tangent space *T*_**x**_ℳ(*t*) has an analytic parametrization, providing an alternative formalism to compute Λ and _*iso*_Λ (SI Sec. 1.1). However, parameterizing dynamic surfaces and using the parametrization to quantify Lagrangian deformation induced by flows on the surface is cumbersome due to singularities of the parametrization. For example, parameterizing a two-sphere 𝒮^2^ using **p** = (*θ, ϕ*) as **r**(**p**) = (sin *θ* cos *ϕ*, sin *θ* sin *ϕ*, cos *θ*) works everywhere except the poles (0, 0, ± 1), where it is not invertible. For a static problem, one can add another parametrization with singularities elsewhere and switch between the two to fully parametrize 𝒮^2^ [24]— i.e. construct an Atlas [55]. However, flows on dynamic manifolds would require a cumbersome construction of several charts [55] and sophisticated numerical schemes to switch between them while integrating trajectories.

To avoid these problems and broaden applications to experimental data, we leverage the discrete representation of manifolds as polygon meshes. Given the velocity field data on a discrete mesh (Fig. 2e), we first compute the flow map 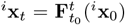 for every initial position ^*i*^**x**_0_ ∈ ℳ(*t*_0_), *i* ∈ {1, …, *N*} where *i* indexes the mesh nodes. We construct local tangent planes and their orthonormal tangent basis vectors [_1_*ζ*(**x**, *t*), _2_*ζ*(**x**, *t*)] on the mesh using the mesh connectivity (SI Sec. 2). Then, we estimate 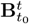 using 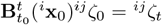 (^*i*^**x**_0_)^*ij*^*ζ*_0_ = ^*ij*^*ζ*_*t*_, where ^*ij*^ *ζ*_0_ = [(^*j*^**x**_0_ − ^*i*^**x**_0_)^⊤^_1_*ζ*(**x**_0_, *t*_0_), (^*j*^**x**_0_ − ^*i*^**x**_0_)^⊤^_2_*ζ*(**x**_0_, *t*_0_)]^⊤^ and ^*ij*^*ζ*_*t*_ = [(^*j*^**x**_*t*_ − ^*i*^**x**_*t*_)^⊤^_1_*ζ*(**x**_*t*_, *t*), (^*j*^**x**_*t*_ − ^*i*^**x**_*t*_)^⊤^_2_*ζ*(**x**_*t*_, *t*)]^⊤^ for every initially nearby node *j* (Fig 2e, SI Sec. 2 for details). From 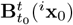, we finally compute 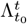 and 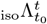.

### Eulerian Coherent Structures

Lagrangian Coherent Structures organize trajectories over any time interval [*t*_0_, *t*]. Given their integrative nature, however, they do not capture short-term variability in material transport. By contrast, Eulerian Coherent Structures [20] and instantaneous Lyapunov exponents [56] provide a rigorous framework to analyze short-time material transport—*short-time attractors* and *short-time repellers*—as well as regions and directions of distinct stretching and shrinking rates. These structures correspond to the instantaneous limit (*t* → *t*_0_) of the corresponding LCS and are computable from the Eulerian rate of strain tensor while remaining inaccessible from Eulerian velocity snapshots [20, 25, 57] and nematic tensor [36]. Specifically, while Lagrangian attractors are marked by ridges of 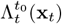 and the associated contracting direction 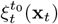, in the limit of *t*_0_ → *t*, short-time attractors are marked by trenches of *s*_1_(**x**, *t*) (i.e. highly negative regions of *s*_1_(**x**, *t*)) and the direction of maximal contraction rate is marked by **e**_**1**_(**x**, *t*), where *s*_1_(**x**, *t*) ≤ *s*_2_(**x**, *t*) are the eigenvalues of the rate of strain tensor and **e**_**1**_(**x**, *t*), **e**_**2**_(**x**, *t*) the correspnding eigenvectors [56]. Similarly, short-time repellers are positive ridges of *s*_2_(**x**, *t*) and **e**_**2**_(**x**, *t*) mark the direction of maximal stretching rate. Here, we extend these techniques to flows on dynamic surfaces (SI Sec. 4).

Lagrangian and Eulerian CSs are complementary. For example, given a velocity dataset over [*t*_0_, *t*_*f*_], computing LCSs over increasing finite time intervals [*t*_0_, *t*_1_], [*t*_0_, *t*_2_] etc., reveals when and where patterns of distinct Lagrangian transport and deformations emerge. By contrast, computing Eulerian Coherent Structures at different *t* ∈ [*t*_0_, *t*_*f*_] reveals the spatiotemporal coordinates of instantaneous high deformation rates. Figure 3a-d and Fig. S4 show Lagrangian attractors and repellers and Eulerian short-time attractors and repellers for the same active nematic system, along with typical velocity and nematic tensor plots.

**Figure 3:**
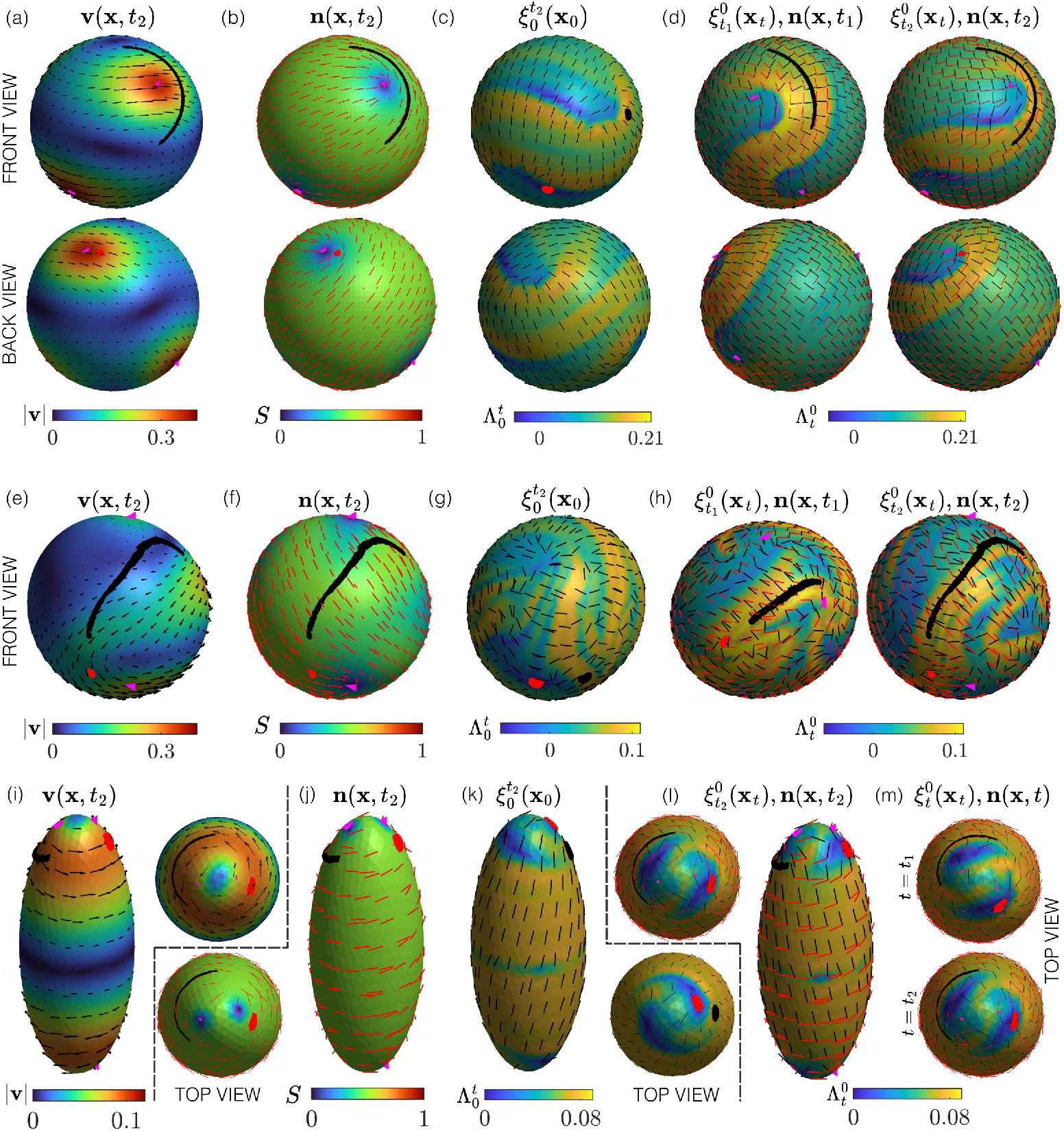
Active nematic flows on stationary and dynamic surfaces : (a-d) Extensile active nematic flow on a stationary spherical surface. a) Velocity field (black arrows) **v**(**x**, *t*_2_) with colormap denoting velocity magnitude. b) Nematic director field (red bars) **n**(**x**, *t*_2_) and nematic order parameter *S*(**x**, *t*_2_) (scalar field). Magenta triangles mark +1*/*2 topological defects. (c) The forward FTLE field 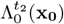 and maximum stretching (or repulsion) direction 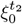 (black bars) for Lagrangian deformation over the time interval [0, *t*_2_]. (d) The backward FTLE field 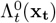, maximum contracting (or attracting) direction 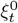 (black bars) and **n**(**x**, *t*) for increasing *t*. a,b,d) Red and black patches mark the final position of tracer particles initialized as small disks at locations shown in (c). SM1 shows the time evolution of the Eulerian and Lagrangian quantities in a-d for different *t*. (e-h) Same analysis as in a-d for an active nematic flow on a deforming vesicle. SM2 shows the time evolutions of panels e-h. (i-m) Same analysis as in a-d for an active nematic flow on a stationary ellipsoid geometry. SM3 shows the time evolutions of panels i-m. All quantities are in nondimensional units.

### 2.2 Active nematic flow on stationary and deformable surfaces

First, we consider an extensile nematic flow on a stationary surface, inspired by experiments with microtubule-kinesin motors confined on a spherical vesicle [10]. While measuring in-toto velocities and nematic directors in these experiments remains challenging, a recent study [58] developed a numerical scheme that recapitulates the flows observed in [10] by solving the nematic hydrodynamic equations on a spherical surface, mechanically coupled to the inner and outer fluids. These equations return the nematic tensor **Q**(**x**, *t*) = *S*(**n** ⊗ **n** − **I***/*2) and the velocity **v**(**x**, *t*) fields, where 0 *< S* ≤ 1 is the nematic order parameter and **n** denotes the mean orientation nematic elements. Because the nematic field is on a sphere, the net topological charge is constrained to +2, which manifests as four +1*/*2 defects. These defects are motile due to activity, and at low activity regimes, the four +1*/*2 defects move along the surface without annihilating each other. Numerical simulations are solved in the small deformation limit of the vesicle surface, which remains approximately fixed.

At low activity, the simulations recapitulate the experimentally observed periodic motion of the four +1*/*2 defects from a tetrahedral to a planar configuration [10, 58], as shown in the first two columns of SM1 and Fig. 3a-b. This periodic motion provides a simple, controllable flow motif to study material transport. Fig. 3c shows the _*f*_ Λ field for the entire time interval [0, *t*_2_], revealing a band-shaped ridge (repeller) in the vesicle at its initial configuration (**x**_0_) where the flow will undergo maximum deformation (or separation) along the 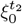 direction normal to the ridge. To visualize this repulsion, we initialize at time 0 tracer particles in a region with high values of _*f*_ Λ (black Fig. 3c) and at a low values for comparison (red Fig. 3c). Fig. 3a-b show the tracers at the final time *t*_2_, displaying how the black patch deforms considerably compared to the red one during [0, *t*_2_]. Owing to its cumulative Lagrangian nature, the band-shaped repeller, organizing material transport and revealing initial regions **x**_0_ and directions of high stretching, cannot be identified from inspection of the Eulerian **v** and **Q**.

Figure 3d shows the backward FTLE 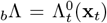 for different *t*. High values of 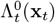 locate the dynamic position (**x**_*t*_) of material patches that undergo maximal convergence during [0, *t*]—i.e. the attractors—along the direction 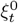. The black patch visualizes this effect, starting initially circular and within the domain of attraction of the attractor (SI Sec. 3 and Fig. S3) and ultimately deforming and converging into the 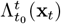 ridge (Fig 3d). Similar to 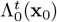 (visualized at the initial configuration), 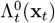 (visualized at the final configuration) reveals a band-shaped structure of the attractor and regions of high contraction, which are not contained in **v** and **Q** plots (Fig 3a-b). Consistent with 2D planar active nematics [36], we find that i) motile +1*/*2 defects deform the _*b*_Λ ridges and ii) the extensile nematic director aligns to the _*b*_Λ ridge (Fig 3d). SM1 shows the time evolution of the Eulerian and Lagrangian quantities in Fig 3a-d. For short-time attractors and repellers associated with Fig. 3a-d, see SI Sec 4 and Fig. S4. This low-activity vesicle system with regular +1*/*2 defects motion offers a simple system to engineer synthetic materials on curved surfaces. The interaction of regular\ +1*/*2 defects motion and the flow attractors and repellers can be leveraged to control material transport and deformation motifs in active nematics on curved surfaces [59–61].

In the high activity limit, the vesicle undergoes both in-the-plane and out-of-plane deformations (Fig. 3e-h), as observed experimentally [10]. Similar to the static nematic vesicle (Fig. 3a-d), ridges of _*f*_ Λ mark regions of high Lagrangian stretching (Fig. 3g), visualized by the distinct deformation of the red and black particle patches (Fig. 3e,f,h), initialized as small circular patches at low and high _*f*_ Λ (Fig. 3g). The red patch undergoes small deformation, while the black patch highly deforms and converges to the attractors marked by 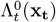 ridges along the 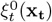 direction (Fig. 3h). Compared to the static vesicle case, Λ ridges display a more complex pattern, indicating higher flow complexity. The director field **n** does not align with the ridges with the _*b*_Λ ridges due to the higher contribution from the elastic interactions in comparison to the low activity limit. SM2 shows the time evolution of the Eulerian and Lagrangian fields associated with Fig. 3e-h.

Recent advances leveraging techniques from discrete differential geometry enabled simulations of nematic-hydrodynamics flows on any Riemannian 2-manifolds [62]. Our approach applies to flows on general surfaces regardless of their genus and curvature variations (Fig 3, Fig. S5). This enables studying the role of curvature, topology, and defect dynamics in the material transport and deformations of active nematics and multicellular flows in deforming organs and organoids. As an example, we study the nematic flows in a prolate ellipsoid, which at low activities result in the periodic motion of two +1*/*2 defects at the poles (Fig 3i-m) [63]. We observe that nematic flow generates a spiral-shaped _*f*_ Λ ridge at the poles with a more spatially homogenous _*f*_ Λ away from the poles (Fig 3k). As before, we visualize this deformation heterogeneity initializing tracer particles at regions of high and low 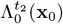 (black, red Fig. 3k), and show their final position in Fig. 3i,j, along with **v**(**x**, *t*_2_) and **Q**(**x**, *t*_2_). Fig. 3l,m show spiral 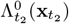 ridges around the poles next to +1*/*2 defects and Fig. 3m shows 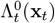 for increasing *t* along with the deformed material patches converging to 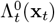 ridges along 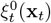 (black patch) or not deforming where 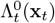 is low (red patch).

### 2.3 Multicellular tissue flows on pancreatic spheroids

Tissue flows in epithelial sheets are ubiquitous in morphogenesis, generating the necessary motion and deformations to form a target shape [2, 64]. We analyze experimental tissue flows on pancreatic organoids [6]—a spherical epithelial monolayer with an apicobasal polarity that undergoes collective cell migration. Tissue velocities are reconstructed using particle image velocimetry (PIV) on live imaging (Fig. 4a and SM4).

**Figure 4:**
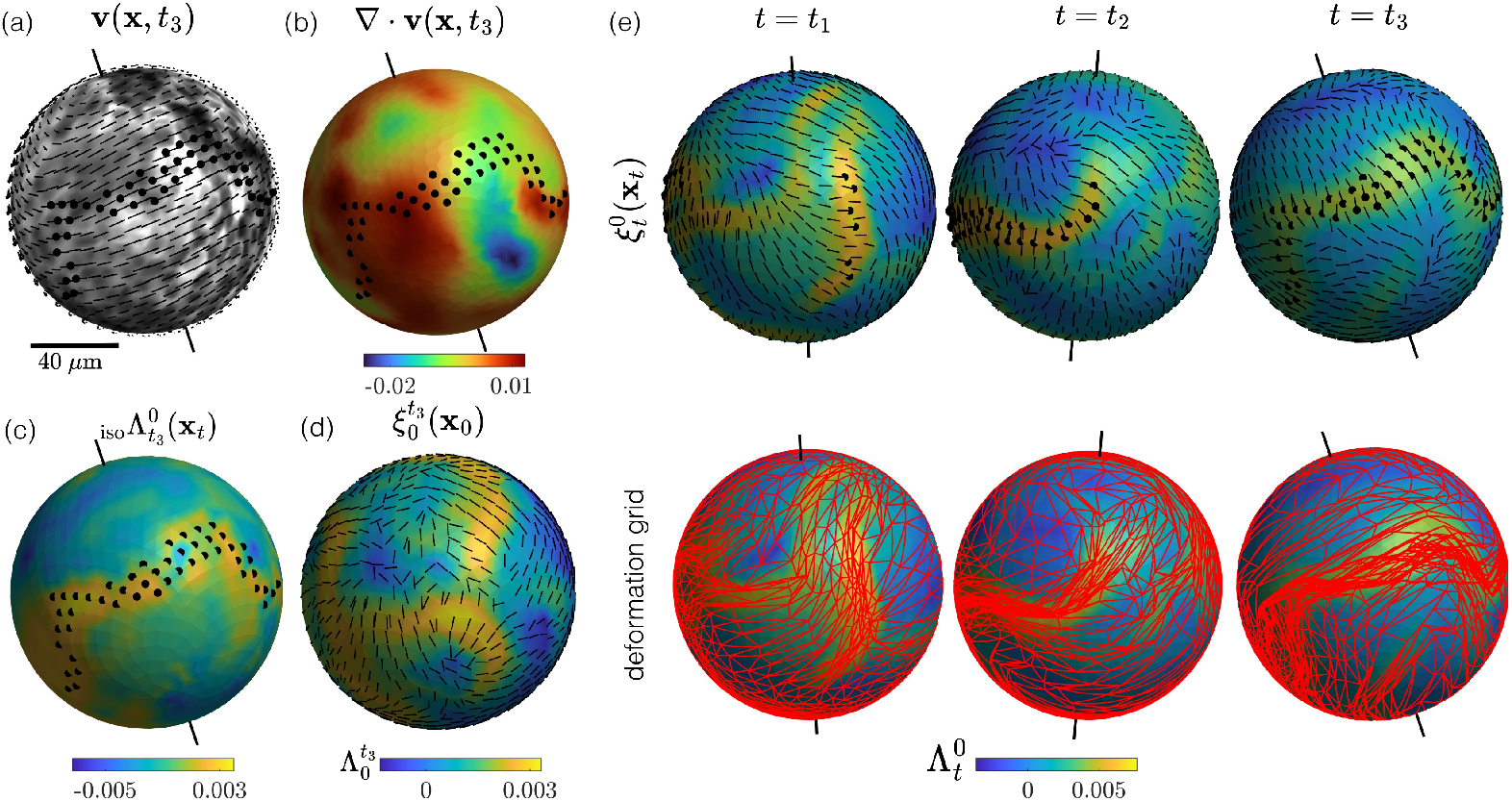
Tissue flows from collective cell migration in pancreatic spheroids: (a) Velocity field overlayed on the apical surface of the spheroid at time *t*_3_ = 330 min. (b) Velocity field divergence ∇ · **v**(**x**, *t*_3_) associated with (a). (c) Isotropic deformation field 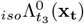. (d) The forward FTLE 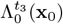 field and the maximum stretching axis 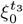 Backward FTLE 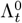 for *t* ∈ {*t*_1_ = 130, *t*_2_ = 230, *t*_3_ = 330} min with the corresponding maximum shrinking direction 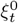 (black bar) in the top row and a deforming Lagrangian grid (red) in the bottom row. (a-e) The instantaneous rotational axis of the spheroid is shown in black (See [6] for details on how the rigid rotation axis is computed from the velocity field). Black dots in a,b,c,e mark regions of high deformation (i.e. regions with 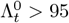 percentile at a given time). SM4 shows the time evolution of panels a-e.

The flow on the surface is compressible (Fig. 4b) generating isotropic Lagrangian deformations, captured by 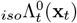 (Fig. 4c).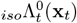 integrates the flow divergence experienced by a material patch along its trajectory, and high 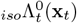 mark final-time locations of tissue patches that experienced large isotropic shrinkage. High values of 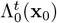 show initial tissue regions that will undergo maximal stretching along 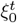, which is perpendicular to the ridges (Fig. 4d). As noted before, 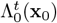 ridges act as a material barrier, whereby tissue patches on either side of this ridge will maximally separate and deform. High values of 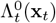, instead, mark the final (**x**_*t*_) position of tissue regions that highly contract along 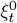 (Fig. 4e, top). These high deformation regions are not contained in Eulerian quantities (cf. high deformation regions are marked by black dots in Fig. 4a,b,c,e and SM4). Fig. 4e bottom shows a deforming Lagrangian grid and 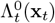 visualizing how grid meshes get attracted to regions of high 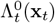 along 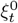. Overall, _*iso*_Λ, Λ and *ξ* fields fully characterize Lagrangian deformations of the pancreatic spheroids.

These material deformation patterns (i.e., felt by the underlying deforming tissue) could instruct different cell responses with implications for cell behaviors and differentiation [17, 65, 66]. Similarly, knowledge of the highest stretching or shrinking axes *ξ* can provide mechanical directional cues to cells, influencing or even generating nematic order and mechanosensitive responses. These deformation patterns captured by LCSs have desirable properties. They are i) computable from kinematic data without requiring the knowledge of the forces generating motion; ii) invariant to the arbitrary choice of the reference systems, hence insensitive to drifts and global rotations; iii) robust to noise, owing to the cumulative nature of LCSs.

### 2.4 Cyclical deformations of a zebrafish embryo heart

Large-scale tissue deformations are ubiquitous for the functioning and morphogenesis of various organs, such as the heart [17, 67], gut [68] and lungs [69]. These cumulative deformations of the tissue can be quantified using Lagrangian methods. We analyze the tissue deformation in a zebrafish heartbeat at 28 hours post-fertilization (hpf) reported in [12]. The zebrafish heart at 28 hpf is a tubular epithelial-like sheet that exhibits cyclical deformations, computable from PIV-derived tissue velocities **v**(**x**, *t*) (Fig 5a, see [12] for details on velocities reconstruction). Computing the forward _*f*_ Λ (Fig. 5b) and backward _*b*_Λ (Fig. 5c) FTLE fields, we observe sharp spatial heterogeneity in the cardiac tissue deformations, not contained in velocity plots (Fig 5a). Spatial heterogeneity in deformations and their temporal dynamics could help uncover regions in the heart with differential material properties, local active stresses, as well as to investigate tissue integrity and mechanosensitive processes [17]. To visualize this deformation, we zoom into a region with high spatial anisotropy of _*b*_Λ field and overlay a deforming Lagrangian grid (Fig. 5c insets). SM6 shows the evolution of the _*b*_Λ field and the deformation grid over time. The ability to identify coherent structures, Lagrangian strains, and principal directions from flows on arbitrary 2D surfaces complements recent computational advances such as TubULAR [12], which quantifies Lagrangian deformation on dynamic tube-like geometries.

**Figure 5:**
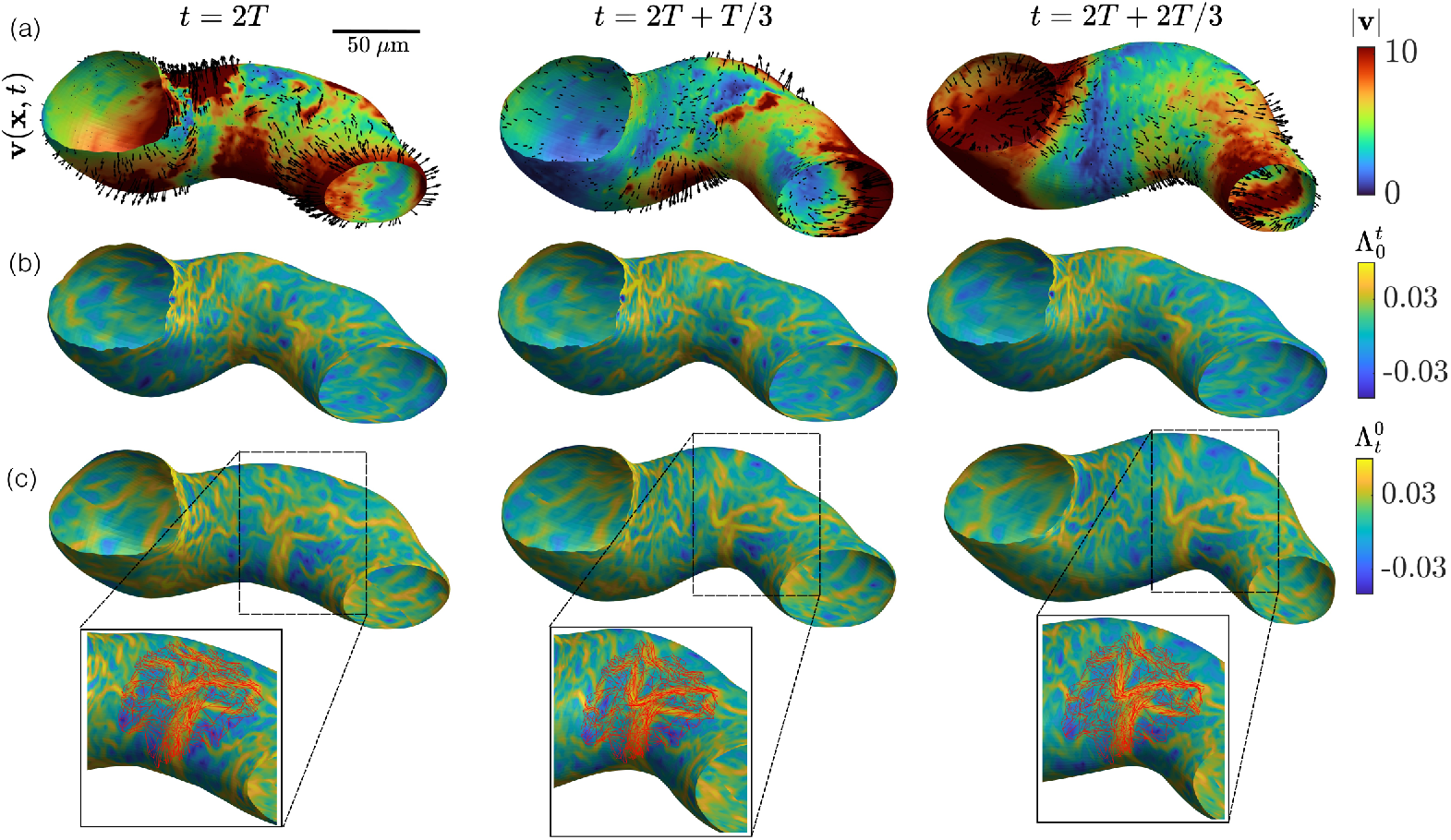
Cyclical deformations of the embryonic zebrafish heart: (a-c) Quantities shown at three different points during a pumping cycle of duration *T*. (a) Velocity field overlaid over the velocity magnitude. (b) The forward FTLE 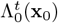 over different values of *t*. (c) The backward FTLE 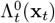 for increasing *t* along with a zoomed-in deforming Lagrangian grid showing contracting deformations on 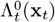 ridges. SM5 shows the time evolution of panels a-b and SM6 of panel c.

## 3 Discussion

Coherent structures organize transport and deformation in steady and unsteady flows, from turbulence and geophysical fluids to plasmas and active matter [18–21, 23–30, 33–39, 41]. Yet, extending this framework to flows on dynamic, curved surfaces—ubiquitous in biological and synthetic systems—has remained unresolved. Here, we developed a geometric, computational framework to identify Lagrangian and Eulerian Coherent Structures (CSs) directly from velocity data on time-evolving surfaces. Our method leverages local tangent plane constructions on triangulated meshes to compute deformation tensors (Fig. 2), revealing anisotropic (Λ, *ξ*) and isotropic (_*iso*_Λ) deformation patterns and dynamic attractors and repellers on evolving manifolds. It generalizes classic LCSs techniques [18, 19, 54] and recent developments in Eulerian CSs [20, 56] to dynamic surfaces with arbitrary curvature, while preserving frame-invariance and robustness to noise.

Applying this framework to active nematic vesicles [10, 58] (Fig. 3, Fig. S4-S5), we uncover Lagrangian attractors and repellers whose spatial patterns and deformation directions are mediated by topological defects and surface geometry. These structures—absent from instantaneous velocity or nematic tensor fields—capture the cumulative effects of activity and geometric frustration, revealing a hidden skeleton of transport that governs how material is stretched, compressed, and segregated. Our results synergize with recent advances linking defect dynamics to flow structures and mixing [35–37], curvature-mediated defect dynamics [11, 60], and optimal control in active systems [70, 71]. These connections highlight emerging opportunities to program material properties by coupling geometry and activity on dynamic surfaces.

Beyond synthetic systems, we demonstrate that collective cell motion in pancreatic spheroids [6] (Fig. 4) and the embryonic zebrafish heart [12] (Fig. 5) exhibits CSs reminiscent of those in active nematics. These biological tissues display dynamically evolving repellers, attractors, and anisotropic deformation axes—kinematic features that might mediate the mechanics of morphogenesis [72], reveal directional cues [65, 73], or regulate mechanosensitive signaling [74–76]. Unlike traditional instantaneous flow diagnostics (e.g., streamlines and vorticity), CSs provide a reduced, frame-invariant description of deforming tissue flows. Thanks to their kinematic nature, CSs can be computed from experimental velocities without knowing the underlying forces.

This framework opens avenues for future exploration. Theoretically, it enables the formulation of inverse problems, such as determining optimal activity or curvature to engineer desired material transport [61, 70, 71, 77]. Experimentally, it invites integration with emerging high-resolution tissue imaging and cell tracking platforms [13–15,42], facilitating inference of mechanical cues from motion measurements. More broadly, by enabling Coherent Structure analysis on dynamic manifolds, this work provides a kinematic characterization of active and living matter, naturally integrating along trajectories local and global mechanisms across scales, with applications from contractile gels [4, 5] to developing organs [3, 12, 17, 67].

## Acknowledgements

We acknowledge Mohammadhossein Firouznia for providing simulated data on nematic vesicles, Tzer Han Tan for providing experimental data on the pancreas spheroid, and Noah P. Mitchell and Dillon J. Cislo for providing experimental data on the beating zebrafish heart. We acknowledge Albert Chen, David Saintiallan and Alex M. Plum for helpful discussions. MS acknowledges support from NSF PHY-2413073 and NSF CAREER PHY-2443851.

## Author Contributions

M.S. designed and supervised the research. S.S. performed research. S.S. and M.S. wrote the paper. C.Z. helped with the numerical implementations and provided the simulated flow data. B. F. provided the Python implementation of the software.

## Code Availability

The tutorial and software used in this work are available in https://sreejithsanthosh.github.io/FTLEhub/.

## Supplementary Information

### S1 Lagrangian deformation of dynamic surfaces

#### S1.1 Case 1. Surfaces defined by a parametrization

Two-dimensional surfaces embedded in three dimensions can be defined by a parametrization function **r**(**p**) ∈ ℝ^3^ where **p** = [_1_*p*, _2_*p*] ∈ ℝ^2^ are the parameters. The surface motion is then prescribed by a three-dimensional velocity field **v**(**r**(**p**), *t*). The dynamic surface ℳ(*t*) is given by the time − *t* position of trajectories of the initial surface points, described by the flow map (Fig S1a)

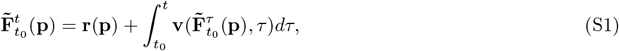

where **r**(**p**) = **x**_0_ parametrizes the surface at initial time ℳ(*t*_0_), and 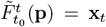 parametrizes ℳ(*t*). To characterize Lagrangian deformations, we parametrize a small neighborhood of **x**_0_ = **r**(**p**), using an infinitesimal vector *δ***p** in the parameter space, which parametrizes the initial material fiber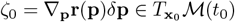. At time *t*, the material fiber is is parametrized by 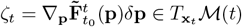 (Fig S1b). The deformation of each material fiber characterizes the local surface deformation. At each surface point labeled by **p**, there is a distinct material fiber that will stretch the most—except for typically isolated locations of isotropic deformations where all fibers stretch equally. The finite-time Lyapunov exponent (FTLE) field 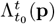 quantifies this Lagrangian maximal deformation rate and is given by

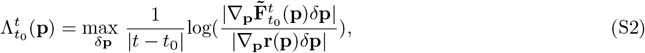

where | · | denotes vector magnitude. Similarly, the Lagrangian isotropic deformation rates of a small neighborhood of **x**_0_ = **r**(**p**) is given by

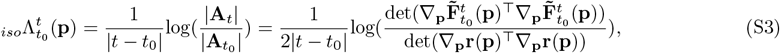

where the the local area element **A**_*t*_ = _1_*ζ*_*t*_ × _2_*ζ*_*t*_, × denotes the cross product and 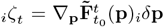. Here, _1_*δ***p**, _2_*δ***p** are two basis vectors spanning the neighborhood of **p** in the parameter space and _1_*ζ*_*t*, 2_*ζ*_*t*_ the corresponding (typically not orthogonal) basis vectors spanning 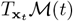.

**Fig. S1:**
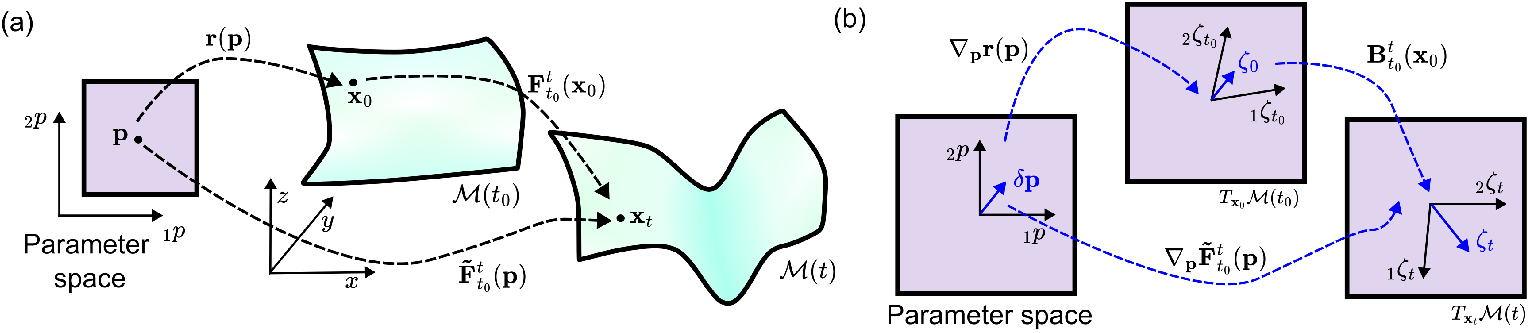
Motion and deformation of a parametrized material surface: (a) The parametrization **x**_0_ = **r**(**p**) maps points **p** in the parameter space to the initial position **x**_0_ ∈ ℳ(*t*_0_). Initial positions will move to later positions according to 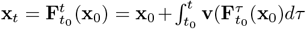. Using **x** = **r**(**p**), one can define 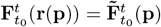 (eq. S1), which parametrizes ℳ(*t*). (b) The matrix ∇_**p**_**r**(**p**) maps infinitesimal vectors *δ***p** in the parameter space to their corresponding material fibers 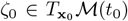. Similary, the matrix 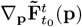 maps *δ***p** to the material fiber 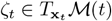.

#### S1.2 Case 2. Non-parametrized surfaces

Typically, the velocity field **v**(**x**, *t*) is provided on a curved surface defined by a discrete set of points (a mesh) without explicit parameterization. Constructing a parameterization and further computing Lagrangian quantities 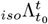 and 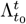 can be cumbersome due to the presence of singularities. For example in a unit-radius two-sphere 𝒮^2^, the parameterization **r**(*θ, ϕ*) = [sin *θ* cos *ϕ*, sin *θ* sin *ϕ*, cos *θ*] generates singularities at *θ* = 0 and *θ* = *π*. At the poles, the parameterization is not invertible, preventing the construction of a basis for the local tangent space and leading to numerical artifacts and instabilities. To overcome this issue, we compute the Lagrangian deformation without a surface parameterization, using the flow map

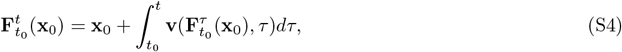

which maps the initial position [*x*_0_, *y*_0_, *z*_0_] = **x**_0_ ∈ ℳ(*t*_0_) to its final position 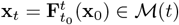 (Fig S1a). To quantify the material surface deformations, we construct an orthonormal tangent vector basis [_1_*ζ*(**x**, *t*), _2_*ζ*(**x**, *t*)] (Sec. S2 for details) for **x** ∈ ℳ (*t*) (Fig. S2). Then, material fibers 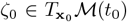 and 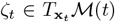 can be represented as

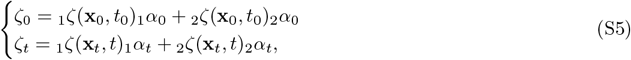

where _*i*_*α*_*t*_ denotes the components of the material fibers in the local tangent space basis. Additionally, the deformation tensor 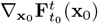 maps *ζ*_0_ to *ζ*_*t*_ by 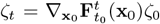. Combining this equation with eq. (S5) gives a linear map 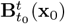 between the components of *ζ*_0_ and *ζ*_*t*_ represented in the local space basis:

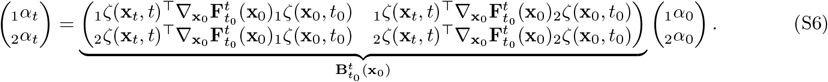

Using the definition of 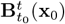 we compute the FTLE field 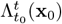 and isotropic deformation 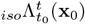 as,

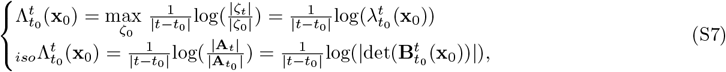

where 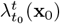 is the largest singular value of 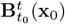. Finally, we note that both 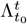 and 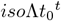 are invariant under arbitrary choices of the tangent space basis used to construct 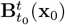.

### S2 Estimating 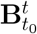 for flows described on a discrete mesh

To compute 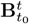, we leverage the representation of curved surfaces as discrete meshes. Flows on a dynamic surface can be described by velocity vectors **v**(^*i*^**x**, *t*), where ^*i*^**x** is the position of the node *i* ∈ {1, ..*N* (*t*)} representing ℳ (*t*) (Fig. S2a). To determine the Lagrangian deformation during [*t*_0_, *t*], we compute the flow map by advecting the initial mesh, i.e., 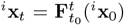, which maps mesh nodes ^*i*^**x**_0_ ∈ ℳ (*t*_0_) at *t*_0_ to ^*i*^**x**_*t*_ ∈ ℳ (*t*). We perform advection using a custom-built Runge-Kutta 4th order method in MATLAB that projects the current position of the mesh node ^*i*^**x**_*t*_ to the surface ℳ(*t*) at each time step. The matrix 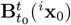 requires constructing the tangent space basis vectors [_1_*ζ*, _2_*ζ*] at ^*i*^**x**_0_ and ^*i*^**x**_*t*_. To construct such basis at a mesh node ^*i*^**x**, we first estimate the normal vector **n**(^*i*^**x**, *t*) by averaging the normal vectors ^*ij*^**n** associated with nearby mesh faces 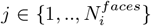 which contains the node ^*i*^**x** (Fig. S2b). Since the normal vectors can either point out or into the surface, we average the tensor quantity 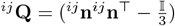 to compute 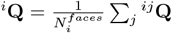. The eigenvector ^*i*^**n** corresponding to the largest eigenvalue of ^*i*^**Q** provides an estimate of the normal vector to the tangent space at ^*i*^**x**. The orthonormal tangent vector basis can then be computed from ^*i*^**n**, by selecting a unit vector _1_*ζ* perpendicular to ^*i*^**n** and _2_*ζ* = _1_*ζ* × ^*i*^**n** (Fig. S2b). Similarly, we compute the tangent vectors at ^*i*^**x**_*t*_ using the normal vector **n** of the mesh face on which it lies (i.e. is projected to) by selecting a unit vector _1_*ζ* perpendicular to it and _2_*ζ* = **n** × _1_*ζ*. Last, the handedness (right or left) of the tangent vector basis need not be consistent between ^*i*^**x**_0_ and ^*i*^**x**_*t*_ as the FTLE 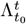 and 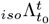 values are invariant to it.

**Fig. S2:**
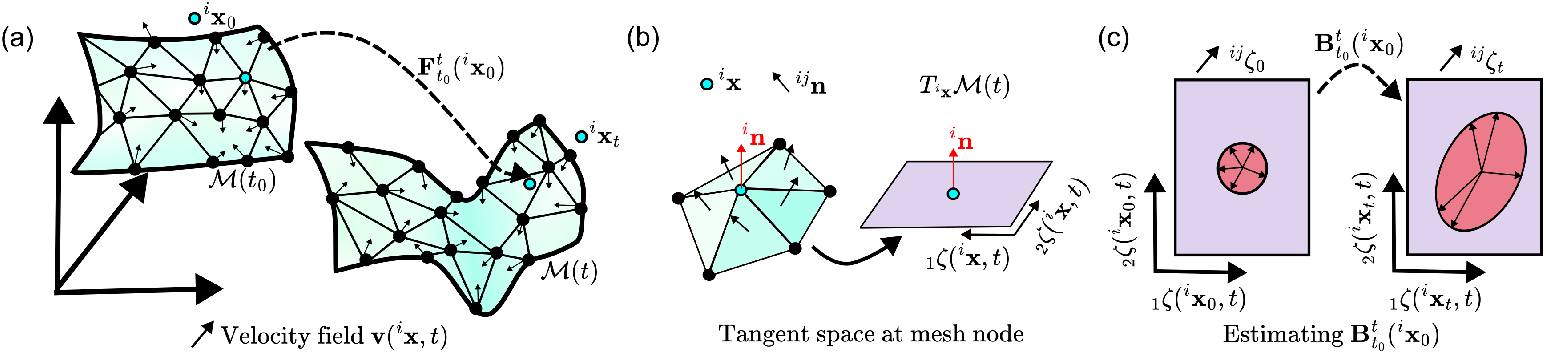
Motion and deformation of a material surface given by a discrete mesh: (a) Flow on curved surfaces are represented by a velocity field **v**(^*i*^**x**, *t*) defined on a discrete mesh ℳ(*t*), where ^*i*^**x** is a mesh node. The flow map 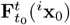 maps the material particle initialized at mesh node ^*i*^**x**_0_ to the final location ^*i*^**x**_*t*_ (typically not at a mesh node). (b) To construct a orthonormal tangent basis [_1_*ζ*(^*i*^**x**, *t*), _2_*ζ*(^*i*^**x**, *t*)] at mesh node ^*i*^**x**, we compute the normal vector ^*i*^**n** by averaging the normal vectors ^*ij*^**n** of nearby mesh faces. (c) 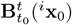 is estimated as the linear mapping between ^*ij*^*ζ*_0_ and ^*ij*^*ζ*_*t*_.

The deformation gradient 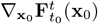 quantifies the variation in the flow map with respect to variations in initial positions **x**_0_. Since the flow is restricted to a surface, the deformation gradient 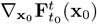 is not defined for variation of **x**_0_ perpendicular to the material surface. Indeed, the matrix 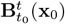 (eq. S6) restricts the variation of the deformation gradient onto the tangent space of the surface. We estimate 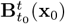 using the relation 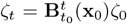 (eq. S5-S6) where 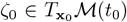 and 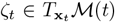 (Fig. S2c). Given the flow map 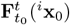, we can define material fibers (^*ij*^*ζ*_0_, ^*ij*^*ζ*_*t*_) in the local tangent vector basis as

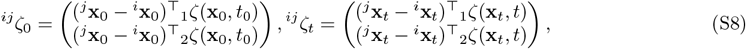

where ^*ij*^*ζ*_0_ is the position vector of neighboring nodes *j* ∈ {1, .., *N*_*i*_} relative to node *i* projected onto the local tangent space at ^*i*^**x**_0_. Similarly, the material fiber ^*ij*^*ζ*_*t*_ is the final position vector of the tracer particle initialized at node *j* relative to the final position vector of the tracer particle initialized at node *i*, projected onto the local tangent space at ^*i*^**x**_*t*_ (Fig. S2c). Because 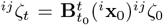 holds for every nearby node *j* ∈ {1, .., *N*_*i*_}, we estimate 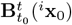 as

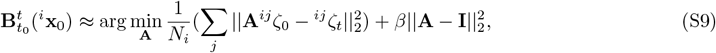

where **A** ∈ ℝ^2×2^, **I** is the identity matrix, *β* is a regularization parameter and || · ||_2_ denotes the Euclidean norm. The term *β* || **A** − **I** ||^2^ regularizes the optimization problem, biasing 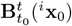 towards identity in the absence of data. This optimization method successfully estimated deformation gradients from sparse and noisy trajectories in 2D and 3D flows [44]. Eq. (S9) can be solved analytically and has the solution

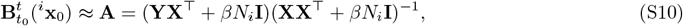

where the matrices 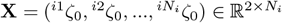 and 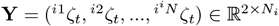.

### S3 Domain of attraction of an attractor

**Fig. S3:**
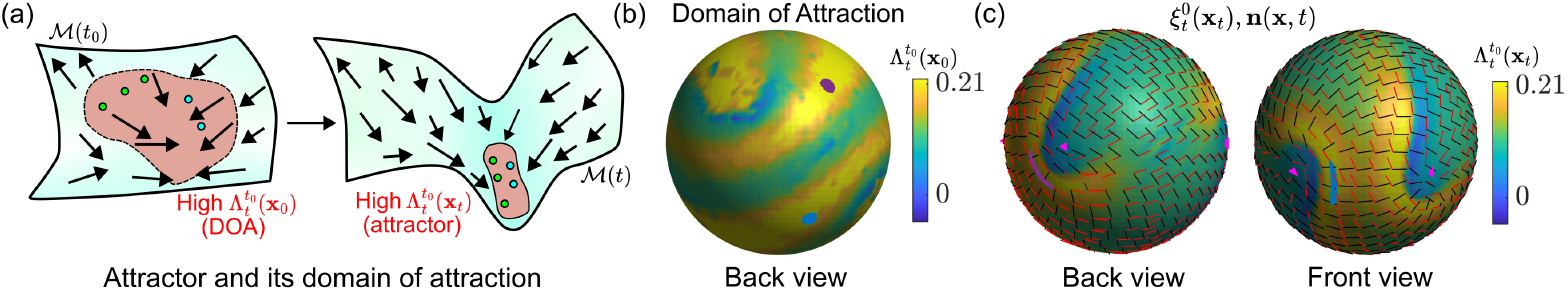
Domain of attraction (DOA): (a) The region with high value of backward FTLE, visualized at the initial configuration 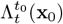 marks the DOA of the corresponding attractor marked by high values of 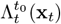 in the final configuration. (b) The DOA visualized using 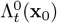 for flow on a static nematic vesicle analyzed in Fig. 3a-d. (c) The backward FTLE 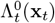, the largest deformation axis 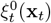 (black bars) and the nematic director field **n**(**x**, *t*) (red bars). (b-c) The violet and blue patches visualize transport starting from different DOAs. The time-evolution of the Eulerian and Lagrangian quantities of the flow shown in (b-c) is given in SM7.

To identify the attracting LCSs (or attractors) in the flow over a time interval [*t*_0_, *t*], we compute the backward FTLE field 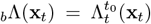 defined on the final position **x**_*t*_. Regions with a high value of _*b*_Λ mark the attractors. In general, there are multiple attractors, each with their corresponding domain of attraction (DOA) at the initial fluid configuration **x**_0_ [38]. The DOA consists of the initial regions (**x**_0_) that will move to the attractor by time *t*, and can be computed by visualizing _*b*_Λ at the **x**_0_, i.e. 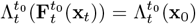 (Fig. S3a). As an example, we visualize the DOA (Fig. S3b) for the flow on an extensile nematic vesicle in the low activity limit, analyzed in Fig. 3a-d. We observe that for each attractor band (Fig. S3c), there is a corresponding domain of attraction. Material patches that start in the DOA (see violet and blue patch, Fig. S3b) deform and move to the corresponding attractors (Fig. S3c).

### S4 Eulerian Coherent Structures on dynamic surfaces

Recent theoretical advances [20] have established frame-invariant criteria for identifying Coherent Structures that influence material transport over short times, providing information undetectable by standard methods [25, 57,78]. These structures include Objective Eulerian Coherent Structures [20] as well as instantaneous Lyapunov exponents [56]. They offer a mathematically rigorous instantaneous limit as *t* → *t*_0_ for Lagrangian Coherent Structures. Eulerian Coherent Structures can be identified from the eigenvalues *s*_1_ ≤ *s*_2_ and eigenvectors [**e**_1_, **e**_2_] of the strain rate tensor **S** = (∇**v** + ∇**v**^⊤^)*/*2, which encode for the instantaneous deformation rates experienced material fibers. Here, we extend these techniques to flows on dynamic surfaces. To formulate the appropriate rate of strain tensor for flows on curved surfaces, we compute the deformation rate of a material fiber *ζ* ∈ *T*_**x**_ ℳ(*t*) following the setup described in Sec. S1.2, which gives

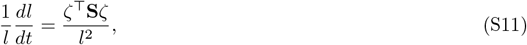

where *l*^2^ = *ζ*^⊤^*ζ* is the fiber length squared. Representing the material fiber in the tangent basis *ζ* = _1_*α*_1_*ζ*(**x**, *t*) + _2_*α*_2_*ζ*(**x**, *t*), we rewrite Eq. (S11) as

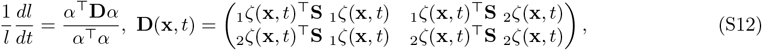

where *α* = [_1_*α*, _2_*α*]^⊤^ and **D**(**x**, *t*) is the rate of strain tensor projected onto the tangent plane *T*_**x**_ℳ(*t*). To estimate **D**(^*i*^**x**, *t*) at mesh node ^*i*^**x** ∈ ℳ(*t*), following the setup described in Sec. S2, we use the relation

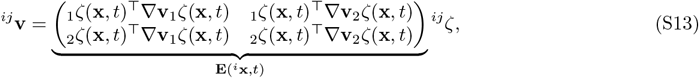

where ^*ij*^**v** = [(**v**(^*j*^**x**, *t*) − **v**(^*i*^**x**, *t*))^⊤^_1_*ζ*(**x**, *t*), (**v**(^*j*^**x**, *t*) − **v**(^*i*^**x**, *t*))^⊤^_2_*ζ*(**x**, *t*)]^⊤, *ij*^*ζ* = [(^*j*^**x** − ^*i*^**x**)^⊤^_1_*ζ*(**x**, *t*), (^*j*^**x** − ^*i*^**x**)^⊤^_2_*ζ*(**x**, *t*)]^⊤^, **D**(^*i*^**x**, *t*) = (**E**(^*i*^**x**, *t*) + **E**(^*i*^**x**, *t*)^⊤^)*/*2 and node *j* and *i* are neighboring nodes. Since Eq. (S13) holds for every neighboring node *j* ∈ {1, .., *N*_*i*_}, we estimate **E**(^*i*^**x**, *t*) as

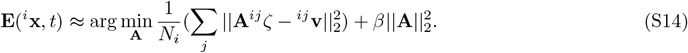

The solution for the optimization given in Eq. (S14) is given by

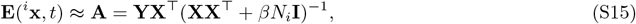

where 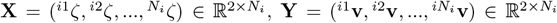 and *β* is a regularization parameter. Using Eq. (S15), we compute **E**(^*i*^**x**, *t*) and its symmetric part **D**(^*i*^**x**, *t*) whose eigenvalues and corresponding eigenvectors are denoted by (*s*_1_ ≤ *s*_2_) and [**e**_1_, **e**_2_]. Regions with high positive values of *s*_2_(**x**, *t*) correspond to short-time repellers, and **e**_2_ marks the direction of repulsion [56]. Similarly, regions of highly negative values of *s*_1_(**x**, *t*) mark short-time attractors and **e**_1_ the direction of attraction.

**Fig. S4:**
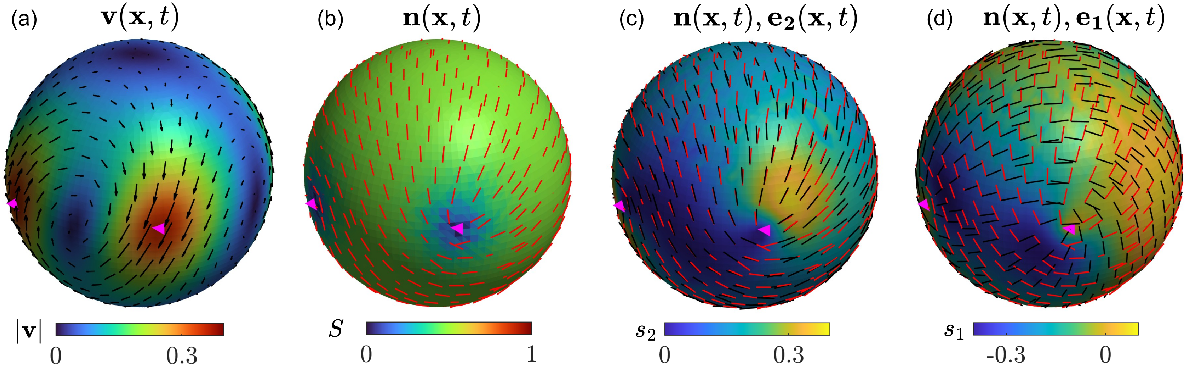
Eulerian Coherent Structures for Flow on a Vesicle (as in Fig. 3a-d): (a) Velocity field with the colormap denoting the velocity magnitude. (b) The nematic director field and the order parameter field. (c) The largest eigenvalue *s*_2_ of the strain rate tensor and corresponding eigenvector **e**_2_ (black arrows) along with the nematic director field (red bars). High values of *s*_2_ mark short-time repellers or regions with the largest stretching rate. (d) The smallest eigenvalue *s*_1_ of the strain rate tensor and the corresponding eigenvector **e**_1_, along with the nematic director field. Highly negative values of *s*_1_ mark short-time attractors or regions with the largest shrinking rate. SM8 shows the time evolution of these quantities.

As a representative example, we compute short-time attractors and repellers (Fig. S4c-d) for the nematic flow on a vesicle analyzed in Fig. 3a-d. Fig. S4c shows a region with the highest stretching rate (high *s*_2_(**x**, *t*)) behind a positive +1/2 defect (magenta triangle), along the **e**_**2**_(**x**, *t*) (black bars), aligned with the nematic director (red bars). Similarly, Fig. S4d shows a region with the highest shrinking rate (highly negative *s*_1_(**x**, *t*)) ahead of a positive +1/2 defect along the **e**_**1**_(**x**, *t*) (black bars), perpendicular to the nematic director. Notably, even Eulerian Coherent Structures remain undetected by inspection of the Eulerian velocity (Fig. S4a) and nematic tensor (Fig. S4b), consistent with earlier observations [20, 36].

### S5 Additional Supplementary Figures

**Fig. S5:**
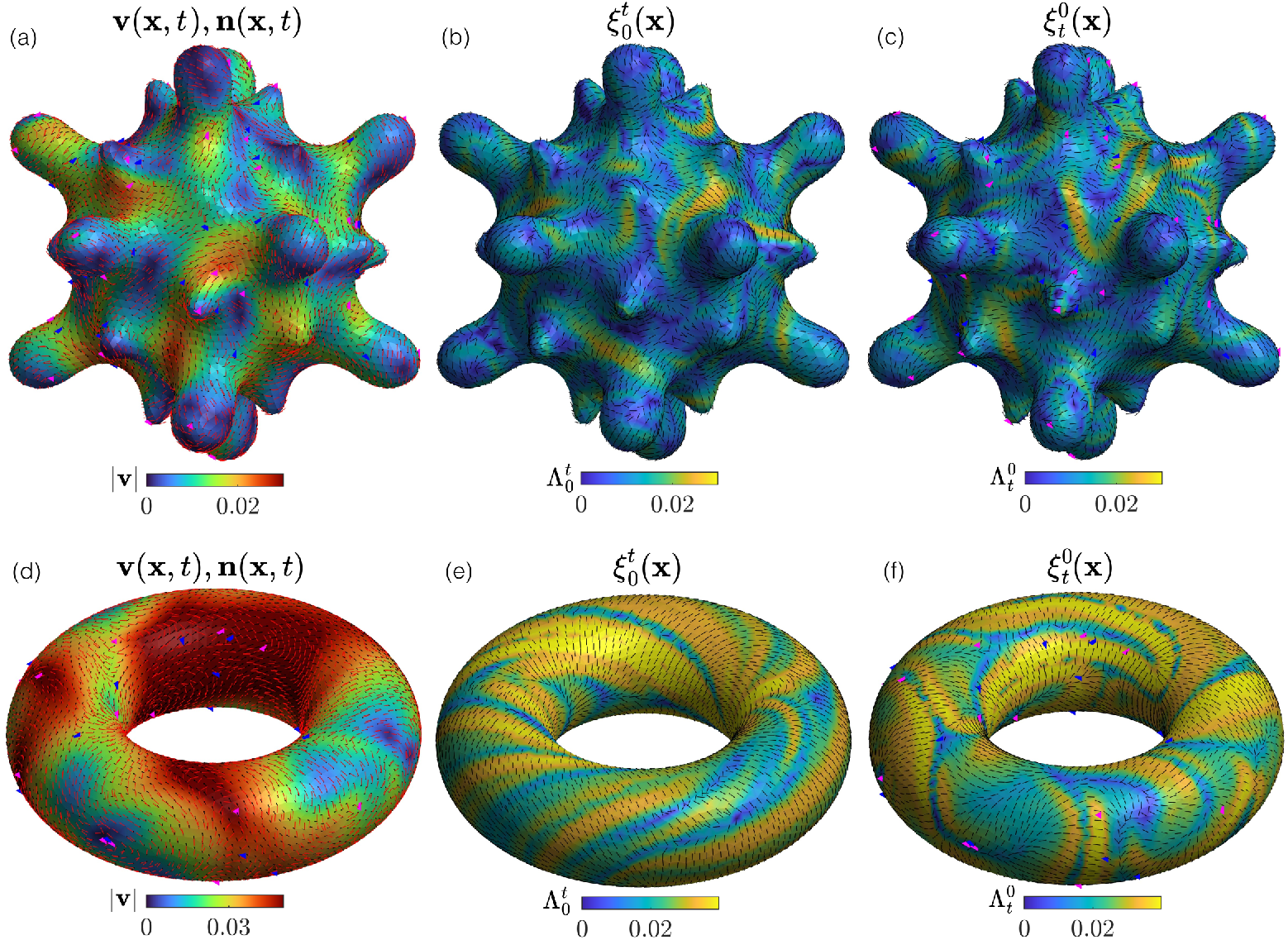
Nematic flow on a topological sphere with heterogeneous curvature and a torus. Nematic flows are simulated using the method described in [62]. Magenta and blue triangles mark the +1*/*2 and − 1*/*2 topological defects. (a) The velocity field **v**(**x**, *t*) (black), nematic director field **n**(**x**, *t*) (red) and the velocity magnitude. (b) The _*f*_ Λ field over the time interval [0, *t*] and the maximum deformation axis 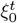. (c) The _*b*_Λ field over the time interval [0, *t*] and maximum deformation axis 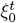. SM9 shows the time evolution of the velocity field 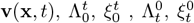 and the nematic director field **n**(**x**, *t*) for varying values of *t*. (d-f) Same as (a-c) but for nematic flow on a torus. SM10 is the same as SM9 but for the surface geometry shown in (d-f).

